# Human dental pulp stem cells: a sanctuary for relapsing *Bartonella quintana*

**DOI:** 10.1101/2020.05.13.094037

**Authors:** Hamadou Oumarou Hama, Attoumani Hamada, Gérard Aboudharam, Éric Ghigo, Michel Drancourt

**Author notes:** These two authors equally contributed to this work. **<u>Corresponding author:</u>** Michel DRANCOURT, IHU Méditerranée Infection, 19-21 Boulevard Jean Moulin, 13385 Marseille, France.

## Abstract

*Bartonella quintana* is a facultative intracellular bacterium responsible for relapsing fever, an example of non-sterilizing immunity. The cellular sanctuary of *B. quintana* in-between febrile relapses remains unknown but repeated detection of *B. quintana* in dental pulp specimens suggested long-term half-life dental pulp stem cells (DPSCs) as candidates. As the capacity of DPSCs to internalize microscopic particles was unknown, we confirmed that DPSCs internalized *B. quintana* bacteria: Gimenez staining and fluorescence microscopy localized *B. quintana* bacteria inside DPSCs and this internalization did not affect the cellular multiplication of DPSCs during a one-month follow-up despite the increase in the bacterial load. *B. quintana*-infected DPSCs did not produce Tumor Necrosis Factor-α whereas an important production of Monocytes Chemoattractant Protein-1 was observed. These unprecedented observations suggested the possibility that DPSCs were shelters for the long-term persistence of *B. quintana* in the host, warranting further experimental and clinical investigations.

## INTRODUCTION

*Bartonella quintana* is a facultative intracellular gram-negative bacterium described in 1915 as the agent of trench fever, emerging during World War I in soldiers presenting with fever, headache, sore muscles, bones and joints and skin lesions on the chest and back^1,2^. Trench fever is nowadays understood as one of the clinical forms of *B. quintana* bacteremia, also responsible for life-threatening endocarditis^3–6^ and bacillary angiomatosis in immunocompromised patients^7^. Further, *B. quintana* is also responsible of lymphadenopathy in the lymphatic territory of its inoculation as *B. quintana* is an ectoparasite-borne pathogen transmitted from person to person by the body lice^1,8^ and probably from cat to persons by cat fleas^9^.

Trench fever is a relapsing fever and *B. quintana* has been consistently observed in circulating erythrocytes during febrile episodes^10^ yet the site where *B. quintana* is residing in-between febrile episodes remains unknown even though the demonstration that bacillary angiomatosis results from the reactivation of quiescent *B. quintana* suggested such a role for endothelial cells as sanctuary cells^11,12^. However, neither erythrocytes nor endothelial cells have been demonstrated to host *B. quintana* for long, in agreement with the fact that both cell types have a limited life span time of 120 days for erythrocytes and 100 days for endothelial cells^13,14^.

Interestingly, *B. quintana* has been consistently detected in the dental pulp, a highly vascularized organ with high erythrocyte trafficking^15^. As for an example, paleomicrobiology studies detected *B. quintana* with a prevalence of 2.5% to 21.4% in the dental pulp collected in buried populations^16,17^. In one particular burial site of Remiremont, the prevalence of *B. quintana* in 45 dental pulp specimens collected from these 5-10^th^ populations, was as high as 53.3%^18^ Also, *B. quintana* has been detected in the dental pulp collected from one patient who had been diagnosed with *B. quintana* bacteremia six months before tooth extraction and was free of bacteremia at the time of tooth extraction^19^.

The dental pulp is composed of several cell types including dental pulp stem cells (DPSCs) which were investigated in the present study. DPSCs are mesenchymal stem cell isolated by Gronthos in 2000 and characterized by the expression of markers such as CD73, CD90 and CD105, whereas markers CD34 (hematopoietic progenitor cell antigen) and CD45 (leukocyte common antigen) are not expressed^20,21^.

The DPSC stemness capacity correlates with a long lifespan^22,23^ making them an attractive cell type to investigate hosting *B. quintana* for extended period compatible with clinical reports. In addition, DPSCs are located in the inner area of dental pulp chamber in close contacts with nerve ending and could be a sentinel cells for injury and blood-borne pathogen invasion. It has been found that DPSCs present an immuno-privileged against immune responses ^24^. Indeed, DPSCs possess an immunomodulatory activity following LPS stimulation. They produce pro-inflammatory cytokines such as Interleukin (IL)-6, IL-8, Tumor Necrosis Factor (TNF)-α and Monocytes Chemoattractant Protein (MCP)-1 to recruit immune cell in the site of inflammation, and anti-inflammatory cytokines including IL-10 to reduce the inflammatory and maintain an homeostasis^25–27^.

Based on this background, the aim of this present study was to investigate the role of DPSCs in host-pathogen interactions, using *B. quintana* as a paradigmal organism.

## MATERIALS AND METHODS

### Bacterial strain

*B. quintana* ATCC49793 was cultured on Columbia 5% sheep blood agar (COS) plates (bioMérieux, Craponne, France) at 37°C under a 5% CO_2_ atmosphere. The identification of *B. quintana* was confirmed by matrix-associated laser desorption ionization/time of flight mass spectrometry (MALDI TOF MS) as previously described^28^.

### DPSCs culture

After obtaining the patient’s informed consent, a wisdom tooth was investigated in line with advice from the IHU Mediterranean Infection Ethics Committee (Advice, 05/29/2018). DPSCs obtained from this wisdom tooth were cultured in Dulbecco’s Modified Eagle Medium F-12 (DMEM/F12, Invitrogen, Villebon-sur-Yvette, France) supplemented with 10% heat-inactivated foetal calf serum (FBS, qualified, EU-approved, South America origin, Gibco, Paisley, UK) at 37°C under a 5% CO_2_ atmosphere. DPSCs viability was determined by using the Trypan blue exclusion assay. This assay is distinctively differentiating non-viable from viable cells based on the analysis of the integrity of the cell membrane^29^. Briefly, 50 μL of trypsinated DPSCs suspension were mixed with 50 μL of a 0.4% solution of Trypan blue dye (Eurobio, Les Ulis, France) for 1 min at room temperature. Cells were immediately counted using a Neubauer microchamber (Brand GmbH, Wertheim, Germany) with a light microscope using a 100 X magnification.

### DPSCs infection

*B. quintana* was collected in sterile tubes from two plates of COS and then washed twice in a row with sterile phosphate buffered saline (PBS). Infection of DPSCs (6 x 10^6^) with *B. quintana* in cell culture medium was performed by centrifugation at 3,220 x g for 1 hour. The suspension was then distributed in flasks (SARSTEDT, Nümbrecht, Germany) (i.e. 2 mL per flask) and incubated at 37°C under 5% CO_2_ atmosphere for a one month follow-up (12h, 24h, 48h, 72h, 1^st^, 2^th^, 3^th^ and 4^th^ week). DPSCs cultured alone and *B. quintana* were used as controls. After each incubation time, cells were washed thrice with sterile PBS and 200 μL of cell suspension were cytospined for 5 min (Shandon Cytospin 4, Thermo Scientific, Cheshire, UK). The identification of infected DPSCs was carried by Gimenez staining.

### Fluorescent *in-situ* hybridization (FISH)

At the second week of infection a FISH was performed after cytospin and fixation of the slides with 4% paraformaldehyde for 20 min at room temperature. FISH was carried out as previously described with some modifications^30^. Briefly, probe 16S488-AATCTTTCTCCCAGAGGG labeled with Alexa-488 fluorochrome (Eurogentec, Angers, France) targeted *B. quintana* 16S rRNA gene. The cellular nucleus was stained in blue using 4’,6-diamidino-2-phenylindole (DAPI, Fisher Scientific, Illkirch, France). Uninfected DPSCs were used as negative controls.

### Cytokine quantification

The supernatants in the first week of the first passage of infected DPSCs and third, fourth weeks of infection were collected to evaluate the concentration of MCP-1 and TNF-α by using ELISA kits according to the manufacturer’s protocols (R&D Systems, Rennes, France), the mean minimum detectable dose of human MCP-1 was 1.7 pg/mL and 4.00 pg/mL for human TNF-α. The results were expressed in pg/mL.

## RESULTS

### Capacity of DPSCs to internalize *B. quintana* bacteria

We first investigated whether DPSCs could internalize *B. quintana* bacteria. Gimenez staining allowing the staining of intracellular bacteria was our reference method. The validation of result was performed using a control of uninfected DPSCs and checking the form of *B. quintana* by Gimenez staining before infection **(Fig. 1)**. *B. quintana* effectively multiplies within DPSCs as indicated by the follow-up from 12 hours of incubation (**Fig. 2**). In addition, FISH was carried out in the second week of infection, confirming the presence of *B. quintana* within cells (in green) **(Fig. 3)**. The specificity of the probe was confirmed by a negative control (**Fig. 4**).

**Figure 1.**
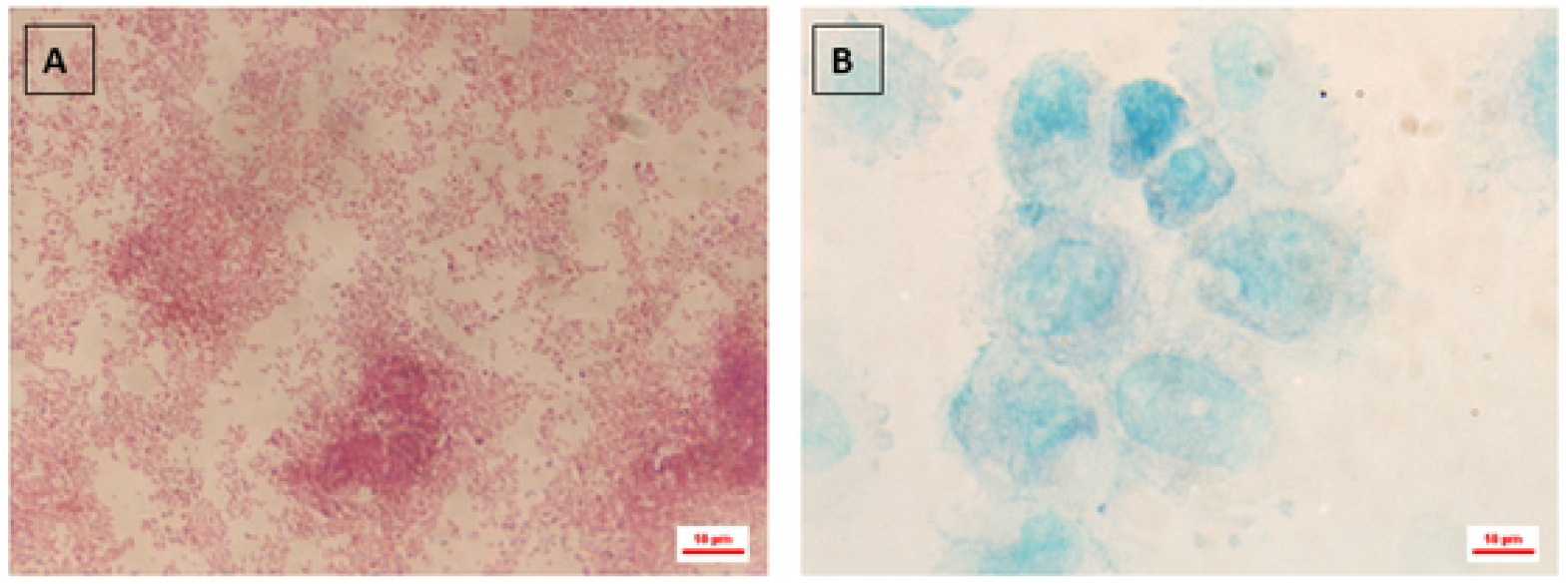
Control of DPSCs and *B. quintana* by Gimenez staining before infection. (A) Form of *B. quintana* colored in red (B): uninfected DPSCs colored in green.

**Figure 2.**
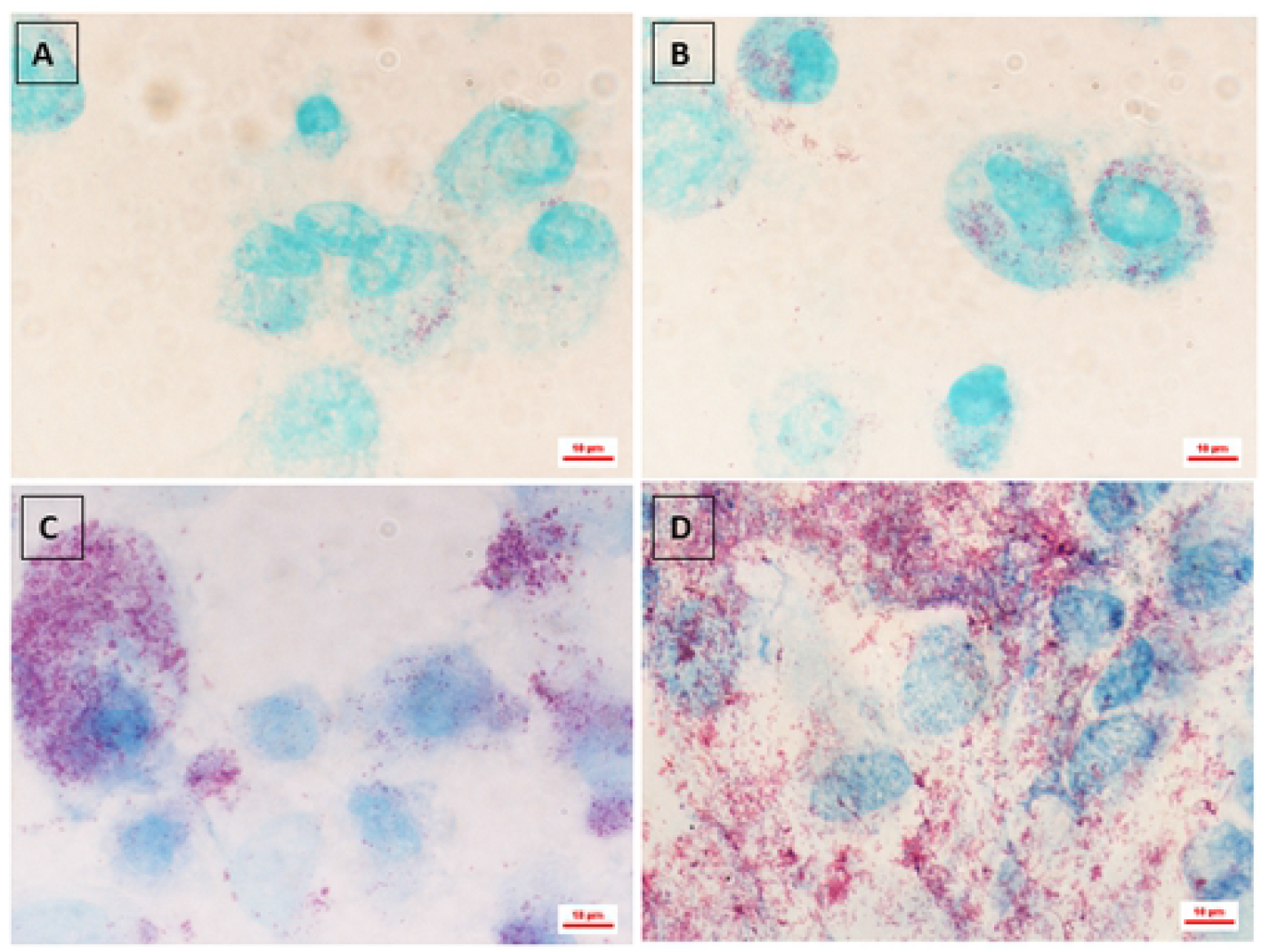
Monitoring DPSCs infection by *Bartonella quintana* (Gimenez staining). (A): 12-hour inoculation (B): 24-hour inoculation (C): One-week inoculation (D): Four-week inoculation. *B. quintana* inside DPSCs.

**Figure 3.**
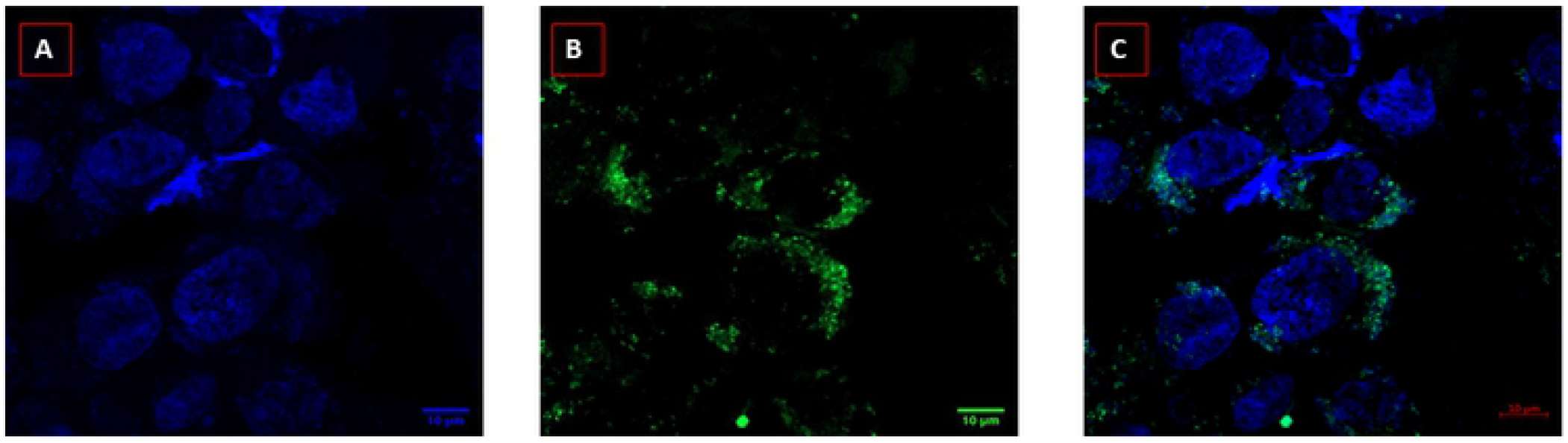
Microscopic FISHing *B. quintana* into DPSCs. (A) DAPI filter (350nm) visualizes cells in blue via the detection of their nuclei (B) FITC filter (488nm) visualizes the 16S rRNA probe in green (C) merge of the two filters (DAPI and FITC).

**Figure 4.**
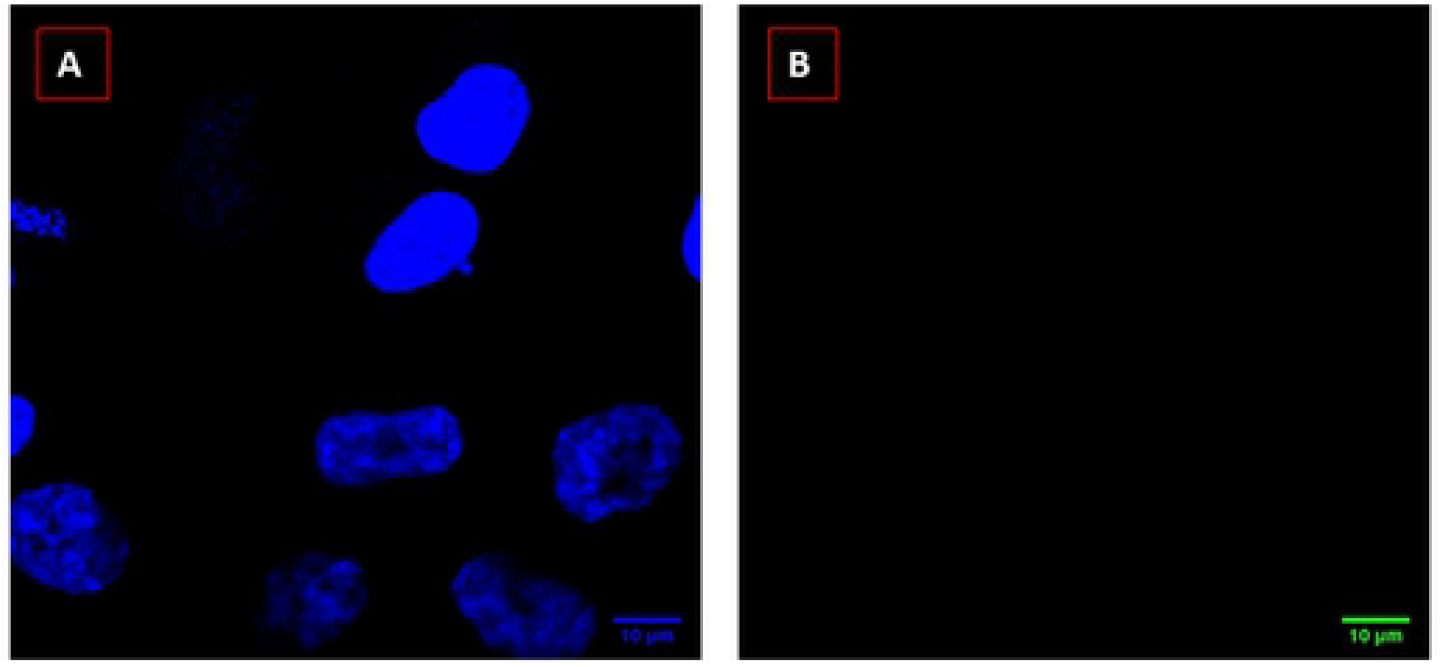
Microscopic FISHing in negative control using the DAPI filter (350 nm) to visualize cells in blue via the detection of their nuclei (A) and FITC filter (488 nm) for the 16S rRNA probe (B).

### Immune response of DPSC to *B. quintana* infection

We further investigated whether *B. quintana* internalization induced an immune pattern by DPSCs. *B. quintana* infection correlated with increased for MCP-1 with maximum of production in week four from 11175 to 19867.85 pg/mL (**Table 1)**. A production of MCP-1 was observed in supernatants of uninfected DPSCs but has a low concentration compared to the infected DPSCs and remains practically stable between the third and the four weeks of incubation (**Table 1)**. For TNF-α, we observed an absence of production suggesting an absence of pro-inflammatory responses TNF-α in infected DPSCs (Supplementary Table S1).

**Table 1.**
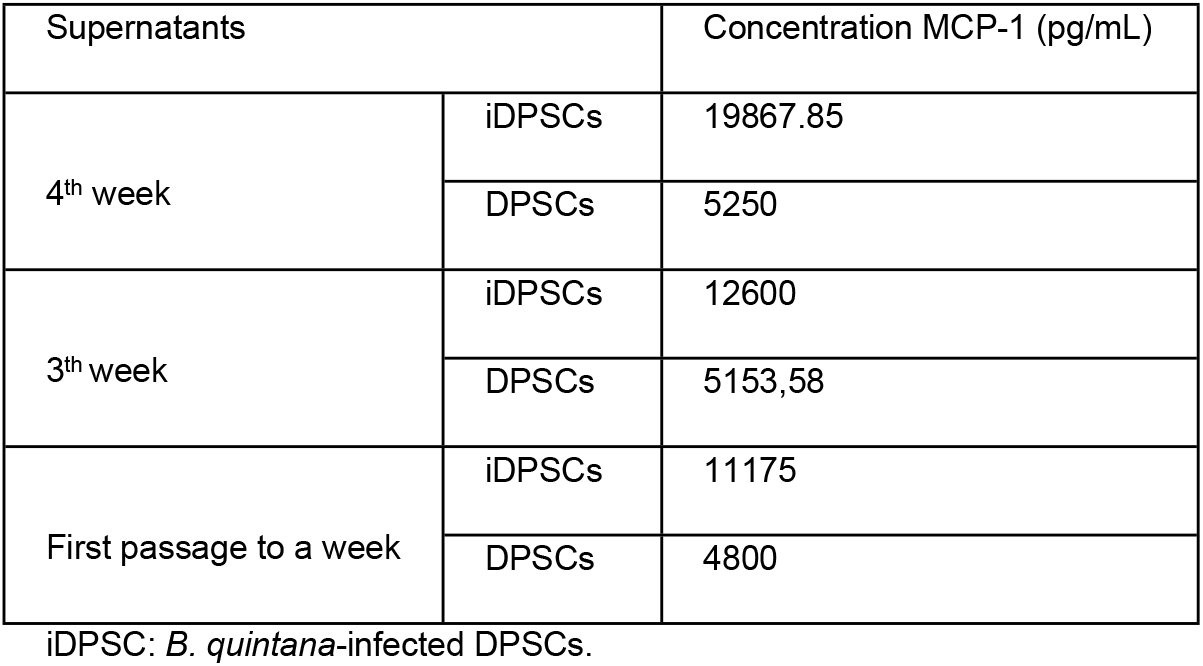
Quantitative determination of human MCP-1 concentrations (pg/mL in cell culture supernatants by ELISA.

## DISCUSSION

We observed that *B. quintana* was internalized by DPSCs. The infection with *B. quintana* did not affected cellular multiplication of DPSCs and despite the increase in these cells an increase in the numbers of bacteria was observed. Our observations support the hypothesis that DPSCs could act as reservoir cells for *B. quintana.* This hypothesis is in accordance with a study reported that intracellular *B. quintana* bacteria could be internalized into a vacuolic compartment (*B. quintana*-containing vacuoles) and multiply^31–33^. These observations correlate with the persistence of *B. quintana* in human^34^. This is opening an exciting new role for DPSCs and further exploring their function could be done by additional observations including the colocalization of *B. quintana* and DPSCs in dental pulp specimens. If confirmed, the role of DPSCs as sanctuary cells for other pathogens will have to be investigated.

The absence TNF-α production suggests that *B. quintana* inhibits or alters the production TNF-α, most likely to favor its replication by avoiding the induction of the microbicidal activities of the DPSCs and the recruitment of pro inflammatory cells. Despite the absence of TNF-α production, production of MCP-1 was observed. This production of MCP-1 has already been described previously as being produced in large quantities by DPSCs^35^ and did not induce DPSCs differentiation according to the literature^36,37^. In addition, it has been described that infection with the Chikungunya virus in human peripheral blood mononuclear cells induce a large production of MCP-1. However, suppression of MCP-1 does not affect replication of virus^38^.

The role of MCP-1 remains unclear in *B. quintana* infection. Further investigations incorporating MCP-1 blocking antibodies may help defining the role of MCP-1 in the replication of *B. quintana* in DPSCs; but this experimental task was beyond the scope of the present study. Nevertheless, this study is opening a new venue for DPSCs as sanctuary cells for the long-term survival of relapsing pathogens.

## ACKNOWLEDGEMENTS

This work was supported by the French Government under the « Investissements d’avenir » (Investments for the Future) program managed by the Agence Nationale de la Recherche (ANR, fr: National Agency for Research), (reference: Méditerranée Infection 10-IAHU-03). This work was supported by Région Provence Alpes Côte d’Azur and European funding FEDER IHUBIOTK.

## AUTHOR CONTRIBUTION STATEMENT

H.O.H. and A.H. performed the experiments, prepared figures. H.O.H., A.H., G.A., E.G. and M.D. designed the experiments, conceived the experiments, analysed the data, wrote the manuscript.

## TABLE LEGEND

**Supplementary Table S1.** Quantitative determination of human TNF-α concentrations (pg/mL) in cell culture supernatants by ELISA.

| Standard curve |  | Samples |  |  |
| --- | --- | --- | --- | --- |
| (pg/mL) | O.D. | Supernatants |  | O.D. |
| 1000 | 1.602 | 4<sup>th</sup> week | iDPSCs | 0.103 |
| 500 | 0.877 |  | DPSCs | 0.101 |
| 250 | 0.538 |  |  |  |
| 125 | 0.354 | 3<sup>th</sup> week | iDPSCs | 0.104 |
| 62.2 | 0.22 |  | DPSCs | 0.099 |
| 31.3 | 0.159 |  |  |  |
| 15.6 | 0.14 | First passage<br>to a week | iDPSCs | 0.105 |
| 0 | 0.1 |  | DPSCs | 0.104 |

342 iDPSC : *B. quintana*-infected DPSCs. O.D. : Optical Density.

